# Wall teichoic acids facilitate the release of toxins from the surface of *Staphylococcus aureus*

**DOI:** 10.1101/2022.01.31.478600

**Authors:** Tarcisio Brignoli, Edward Douglas, Seána Duggan, Olayemi Grace Fagunloye, Rajan Adhikari, M. Javad Aman, Ruth C. Massey

## Abstract

A major feature of the pathogenicity of *Staphylococcus aureus* is its ability to secrete cytolytic toxins. This process involves the translocation of the toxins from the cytoplasm, through the bacterial membrane and the cell wall to the external environment. The process of their movement through the membrane is relatively well defined, involving both general and toxin-specific secretory systems. Movement of the toxins through the cell wall was considered to involve the passive diffusion of the proteins through the porous cell wall structures, however, recent work suggests that this is more complex, and here we demonstrate a role for the wall teichoic acids (WTA) in this process. Utilising a genome-wide association approach we identified a polymorphism in the locus encoding the WTA biosynthetic machinery as associated with the cytolytic activity of the bacteria. We verified this association using an isogenic mutant set and found that WTA is required for the release of several cytolytic toxins from the bacterial cells. We show this effect is mediated by a change in the electrostatic charge across the cell envelope that results from the loss of WTA. As a major target for the development of novel therapeutics, it is important that we fully understand the entire process of cytolytic toxin production and release. These findings open up a new aspect to this process that requires in-depth investigation, while also demonstrating that clinical isolates can utilise WTA production to vary their cytotoxicity, thereby altering their pathogenic capabilities.

**Importance:** The production and release of cytolytic toxins is a critical aspect to the pathogenicity of many bacterial pathogens. In this study we demonstrate a role for wall teichoic acids, molecules that are anchored to the peptidoglycan of the bacterial cell wall, in the release of toxins from S. aueus cells into the extracellular environment. Our findings suggest this effect is mediated by a gradient of electrostatic charge the presence of the negatively charged WTA molecules create across the cell envelope. This work brings an entirely new aspect to our understanding of the cytotoxicity of *S. aureus* and demonstrates a further means by which this major human pathogen can adapt its pathogenic capabilities.

## Introduction

*S. aureus* is a Gram positive human pathogen that can cause a wide array of diseases, ranging from skin and soft tissue infections to bloodstream infections (1, 2). The ability to cause such a wide range of infections is determined by the impressive array of virulence factors that *S. aureus* is able to produce, which include adhesins, immune evasion factors and cytolytic toxins (3–5). The cytolytic toxins produced by *S. aureus* can be classified as receptor mediated toxins and non-receptor mediated toxins (6). Alpha hemolysin (Hla), bi-component toxins like leukocidins (e.g. LukSF, LukAB and LukED) or gamma hemolysin (HlgAB and HlgCB) are receptor dependent toxins, and form stable multimeric pores upon interaction with a specific receptor on target cells (7, 8). Phenol-soluble modulins (PSMs) instead are small peptides that have detergent-like properties, with a broader array of cellular targets (9, 10).

*S. aureus* toxins generally carry out their function in the extracellular milieu, hence they need to cross the bacterial envelope in order to reach this localization. As with most Gram positive bacteria, the *S. aureus* cell envelope is composed of a cytoplasmic cell membrane and a peptidoglycan cell wall (11). Most proteins are translocated through the cytoplasmic membrane by the Sec system, although some proteins are translocated with specialized systems such as the SecA2/SecY2, Tat or Type VII secretion system (12, 13). A further system, named Pmt, is specifically used to translocate PSMs (14). Contrary to the cytoplasmic membrane, the cell wall is a porous polymer and secreted proteins are thought to passively diffuse through it (15–17). However, chemical and physical modifications can influence cell wall architecture and permeability, and the regulation of protein translocation through the cell wall is still an understudied process.

Teichoic acids are a major class of surface polymers found in Gram positive bacteria, which are named wall teichoic acids (WTA) if anchored to the peptidoglycan, or lipoteichoic acids (LTA) if bound to the cytoplasmic membrane (18, 19). These phosphate rich polymers create what has been described as a “continuum of negative charge” that extends from the bacterial membrane to the outermost layers of peptidoglycan (20). LTA and WTA share some functions, and are involved in cell division, cell wall maintenance and turnover (21–24). Despite their similarity, the biosynthesis of these polymers is performed by two distinct pathways. WTA biosynthesis is initiated by the TarO enzyme with the attachment of GlcNAc onto a lipid carrier (25, 26). WTA biosynthesis then continues in the inner leaflet of the cytoplasmic membrane with the contribution of several Tar enzymes, and the polymer is then flipped outward and attached to the peptidoglycan. In contrast, LTA polymerization happens on the outer leaflet of the cytoplasmatic membrane and involves the enzymes encoded by the *ltaA, ltaS* and *ypfP* genes (27, 28). Recently LTA has been associated with the secretion and sorting of the LukAB toxin (29). Differently to the other bi-component toxins, LukAB is partially secreted and partially retained on the cell surface, due to a sorting process that involves the cell membrane lipid lysyl-phosphatidylglycerol (LPG) and LTA. This demonstrates that toxin secretion can rely on complex mechanisms, involving many diverse cell envelope macromolecules.

In previous work, a genome-wide association study (GWAS) identified a locus responsible for WTA synthesis as associated with changes in the cytolytic capacity of *S. aureus* (30). Here we explore the role of WTA in *S. aureus* toxicity. We found that a reduction in WTA production causes a reduction in the secretion of toxins, and an increased abundance of toxins being retained with the cell. This mechanism did not involve a change in cell wall porosity, but instead the toxins appear to be retained due to electrostatic interactions with membrane bound molecules such as LTA. Our results suggest that toxin diffusion through the cell wall is a complex, stepwise mechanism that results from the interplay of a wide range of cell envelope components, and includes the electrostatic gradient that spans from the membrane to the extracellular environment.

## RESULTS

### Clinical isolates with a SNP within the *tarF* promoter region produce more WTA and more cytolytic toxins

In previous work on a collection of 90 clinical MRSA isolates, we identified an association between a single nucleotide polymorphism (SNP) in an intergenic region within the *S. aureus* WTA biosynthetic locus and cytolytic activity. To understand this association, we have ranked the clinical isolates from most to least toxic and indicated which contained this SNP (Fig. 1a). Despite cytolytic activity being a polygenic trait, there is a clear association between the strains containing the SNP and high levels of cytolytic activity. To examine whether this intergenic *tar* SNP affected WTA production, we quantified the phosphate levels in WTA preparations from 10 clinical isolates that contained the SNP and 10 isolates with the reference sequence (i.e. no SNP), and found the SNP containing strains had significantly higher levels of WTA within the bacterial cell wall (Fig. 1b). The SNP is located between the *tarK* and *tarF* genes and is just upstream of the *tarF*-35 site (Fig. 1c). In an attempt to determine the effect the SNP has on the transcription of the *tarF* gene we performed qRT-PCR on the clinical strains, but we were unable to detect any gene transcription. We also cloned this promoter region upstream of a GFP reporter system and detected no fluorescence with either the reference or SNP containing promoter region (Supp. Fig. 1). This suggests that *tarF* is transcribed by these clinical isolates at levels undetectable using these technologies.

**Fig. 1:**
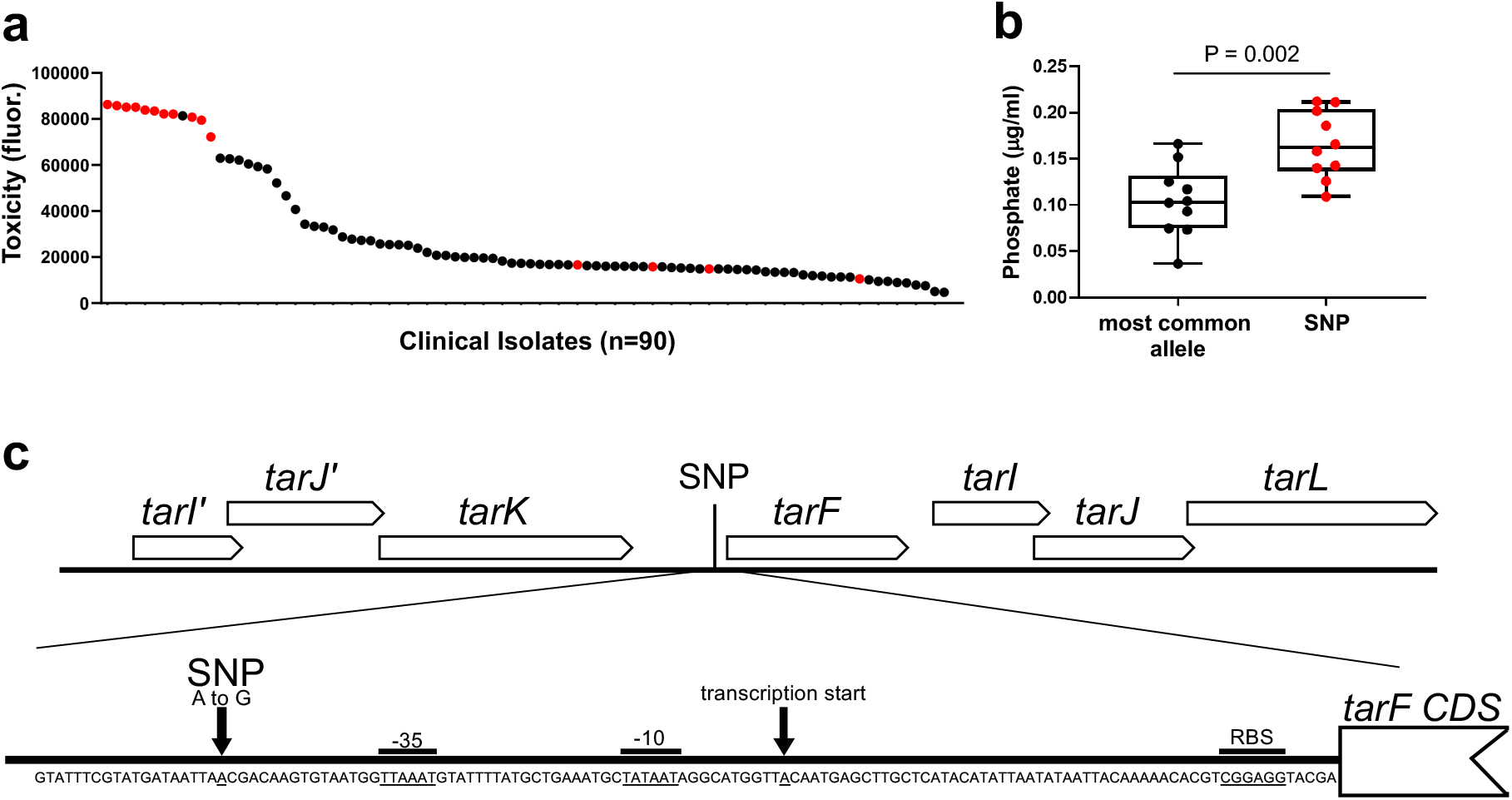
A SNP in the *tar* locus of clinical *S. aureus* isolates associates with higher levels of WTA and cytolytic activity. (**a**) The cytotoxicity of each clinical isolate is presented from highest to lowest with the isolates containing the SNP in the *tar* locus indicated in red. (**b**) The SNP containing isolates produce significantly more WTA compared to an equivalent number of isolates that do not contain the SNP, as measured by phosphate levels in WTA extracts. Error bars represent standard deviation, statistical significance was determined by t test. (**c**) Illustration of the position of the SNP within the Tar locus, promoter features were predicted using Softberry BPROM (softberry.com) (31).

### WTA production affects the cytolytic activity of *S. aureus*

Although we were unable to verify a direct effect of the SNP, our analysis of clinical isolates suggests that mutations that affect WTA production affects the cytolytic activity of *S. aureus.* To examine this further we used a set of isogenic mutants of the MRSA strain LAC, one with low levels of WTA relative to the wild type strain (with transposon insertion in the intergenic region between the *tarK* and *tarF* genes) and one with no WTA (where the *tarO* gene has been deleted). It is worth noting that many of the *tar* genes, including *tarF*, cannot be inactivated due to the toxic effect a build-up of the partially formed components of WTA has on the bacteria (32, 33). This set of strains were selected as it allowed us examine the effect a range of WTA production levels (i.e. high (LAC), intermediate (LAC *tarKF::Tn*) and none (LAC *DtarO*)) had on the cytolytic activity of the bacteria. To demonstrate the effect the mutations had on WTA production we extracted WTA from the cells, visualised the WTA on a tricine acrylamide gel with Alcian blue staining, and quantified the level of WTA production using ImageJ (Fig. 2a & b).

**Fig. 2:**
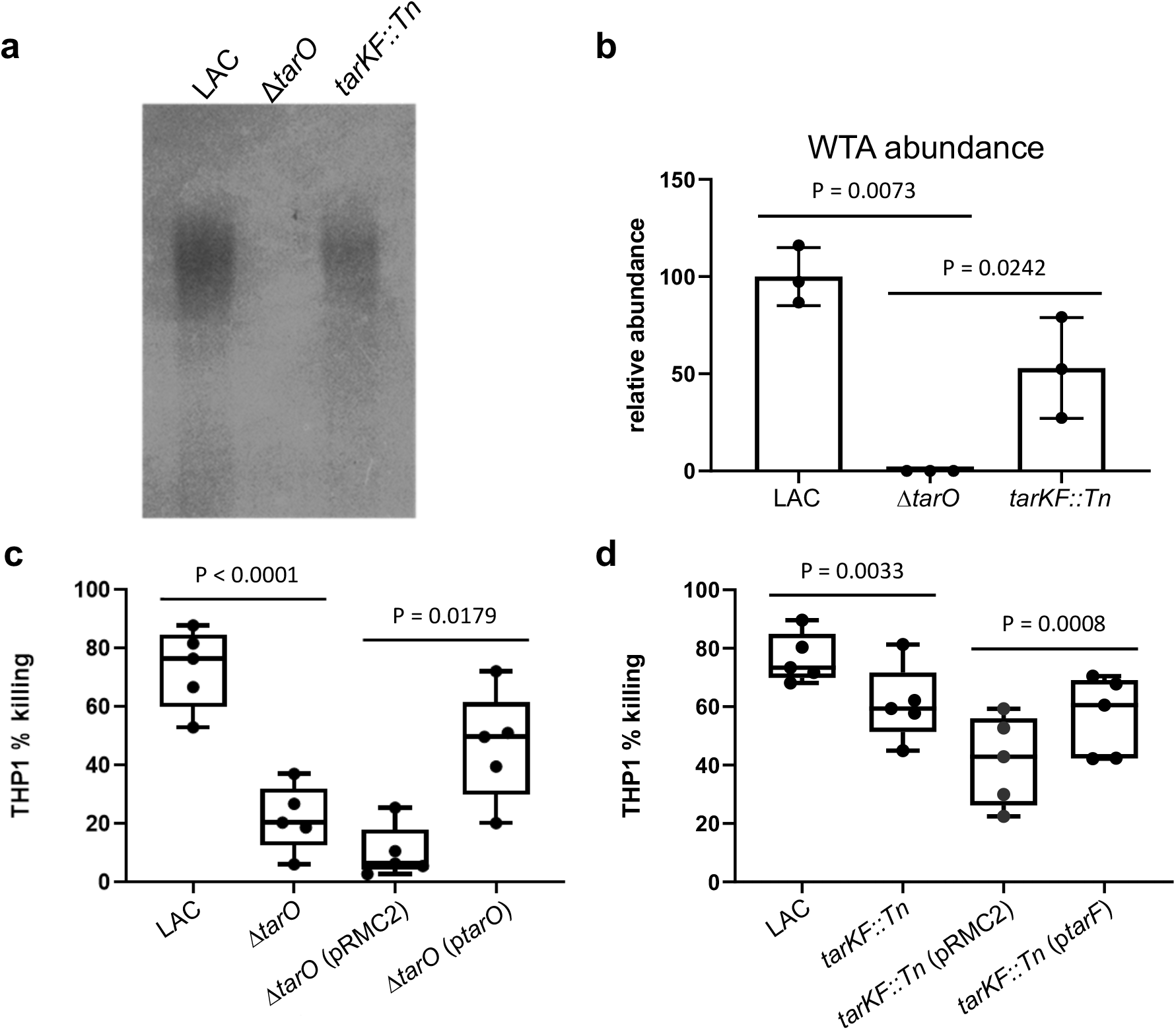
WTA production affects the cytolytic activity of *S. aureus*. (**a & b**) WTA gels and ImageJ quantification, verifiying the differing levels of WTA produced by the wild type LAC strains and isogenic mutant of it with no WTA (LAC Δ*tarO*) and intermediate WTA (LAC *tarFK::Tn*). Error bars represent standard deviation, statistical significance was determined by unpaired t test (**c & d**) THP-1 cell lysis by the isogenic mutant set demonstrating that lower levels of WTA production affects the cytolytic activity of *S. aureus*. This effect was complemented by expressing the respective genes from the pRMC2 plasmid. Each dot represents one biological replicate, error bars represent standard deviation, statistical significance was determined by paired t test.

To verify the effect varying levels of WTA production had on cytolytic activity we incubated bacterial supernatant with cultured THP-1 cells, which are an immortalised monocyte progenitor cells line that are susceptible to the majority of cytolytic toxins produced by *S. aureus* (34). Both mutants were significantly less cytolytic compared to the wild type strain in a manner that was complemented by expressing the respective *tar* gene from the inducible plasmid pRMC2 (Fig. 2c & d). The effect on cytolytic activity was greater in the strain producing no WTA compared to the strain producing an intermediate level, suggesting there is a dose dependent WTA effect on the cytolytic activity of *S. aureus*.

### WTA production affects toxin localisation and secretion

To determine which toxins are differentially produced in the abscence of WTA we performed a series of bacterial supernatant and whole cell lysate extractions and Western blots to quantify the level of cytolytic toxins produced by the bacteria. There was a range of effects on the toxins detected: the AB leukotoxin (LukAB), gamma haemolysin (HlgA) and alpha toxin (Hla) were less abundant in the bacterial supernatants of the WTA mutants compared to the wild type (Fig. 3a), which explains the reduced cytolytic activity of these mutants. However, in the whole cell lysates of the WTA mutants Hla, LukS and LukF were more abundant than in the wild type strain, while LukAB was less abundant and HlgA undetectable in both strains (Fig. 3b). This suggests that the release of some of these toxins from the bacterial cells is less efficient in the WTA mutants. We also examined the location of the phenol soluble modulins (PSMs). These small surfactant-like toxins that migrate ahead of the dye front as a single band of approximately 2-3 kDa on SDS-PAGE gels, which we have verified by mass-spectroscopy (Supp. Table 1). A similar apparent trapping of the PSMs inside the cells was observed (Fig. 3c & d).

**Fig. 3:**
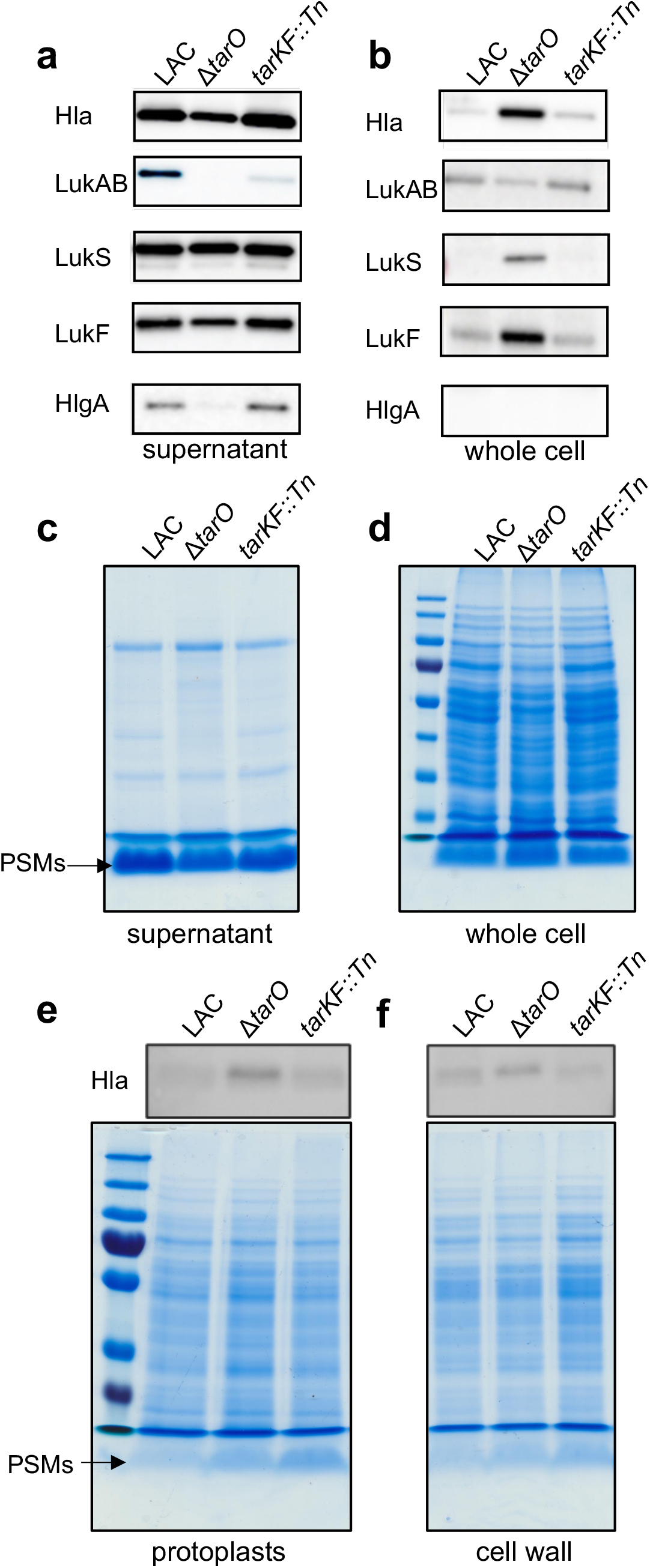
WTA production affects the release of toxins form the S. aureus cell. (**a & b**) Western blots detecting toxins abundance in both intra- and extracellular extracts of the isogenic set of WTA producing strains demonstrating that some toxins are not released from the bacterial cells. (**c & d**) PSM production of the isogenic set of WTA producing strains demonstrating that some of the PSMs are not released from the bacterial cells. (**e & f**) Western blot and PSM gels detecting the abundance of both Hla and the PSM showinng that they are trapped within both the cell wall and protoplast of the *S. aureus* cells. Full length gels and westerns with replicates can be found in Supp. Fig. 2.

To further examine where the toxins are being trapped within the bacterial cells, we separated the cell walls from the protoplasts with a focus on just Hla and PSM abundance, as these were the toxins with the clearest difference in their release from the bacterial cell between the wild type and WTA mutants, i.e. they were more abundant in the whole cell extract of the mutants and less in the supernatant. In doing so we found an increase in abundance of both Hla and the PSMs in both cellular compartments (Fig. 3e & f).

### Passive diffusion of toxin-sized molecules through the cell wall is not impaired in the absence of WTA

In previous studies the inactivation of TarO is reported to have a major effect on the architecture of the cell envelope (21, 35, 36). We confirmed this by transmission electron microscopy, where we also see major morphological changes associated when no WTA is produced, but also an increase in cell wall thickness when there is an intermediate level of WTA produced relative to the wild type strain (Fig. 4a). We therefore hypothesised that passive diffusion of the cytolytic toxins through the cell wall may be inhibited by the different morphology of cell wall of the WTA mutants. To examine whether WTA affects diffusion through the cell wall, we isolated murein sacculi of the wild type and mutant strains, incubated these overnight with fluorescently labelled 40 kDa dextrans and measured the rate of diffusion of the dextrans from the sacculi over a period of 30 minutes. The rate of diffusion of the dextrans was equivalent across the strains, suggesting that the changes to the thickness of the cell wall in the absence of WTA is unlikely to be the explanation for the increase in the amount of toxins trapped within the cell wall of the WTA mutants (Fig. 4b).

**Fig. 4:**
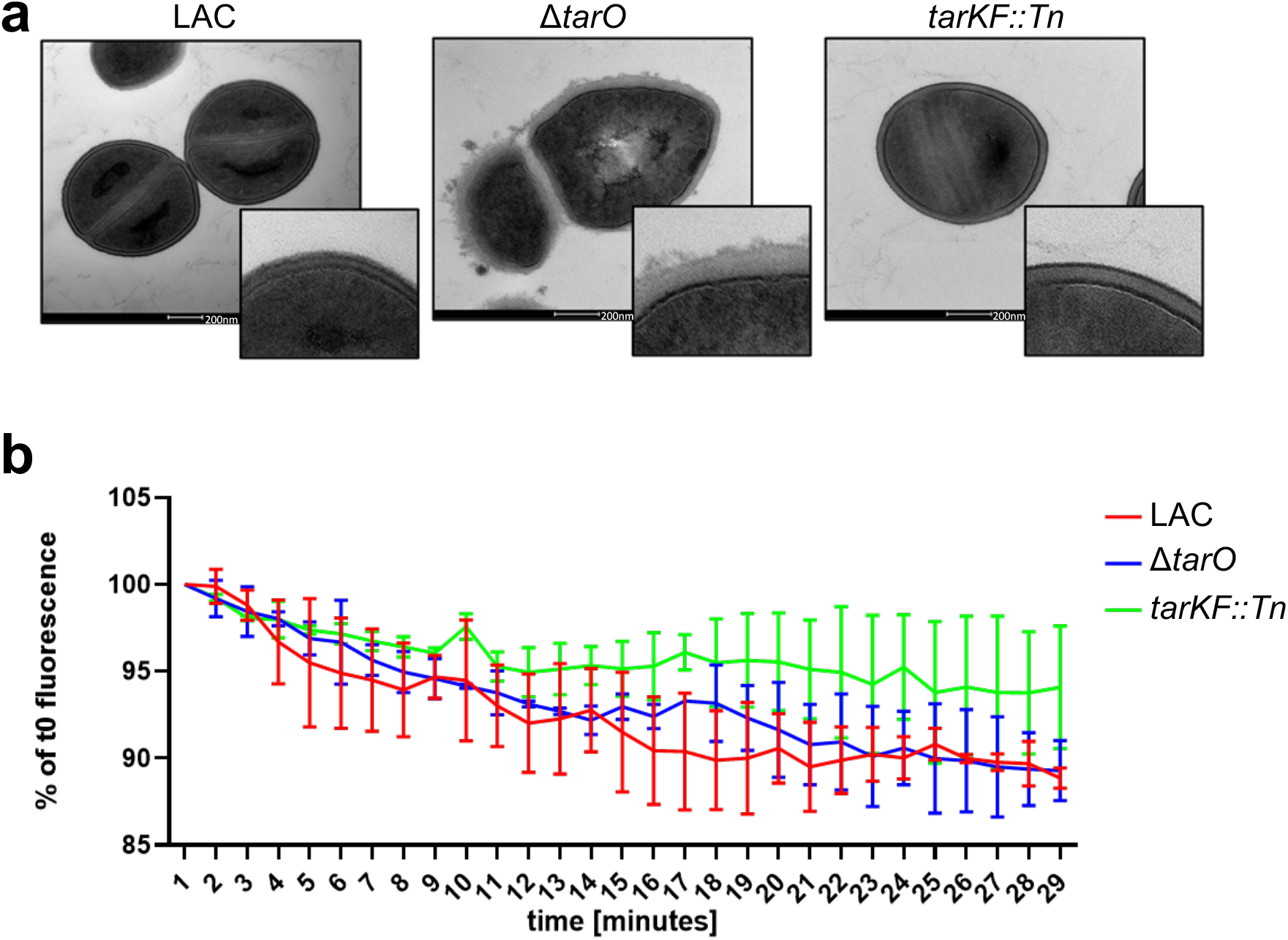
Alterations in the architecture of the cell wall in the abscence of WTA does not affect diffusion through it. (**a**) Transmission electron microcopy images demonstrating the changes differing levels of WTA production make to the *S. aureus* cell wall. (**b**) WTA production levels do not affect dextran diffusion from the murein sacculi of the *S. aureus* strains producing varying amounts WTA. The graph represents two independent biological repeats, bars represent standard deviation.

**Fig. 5:**
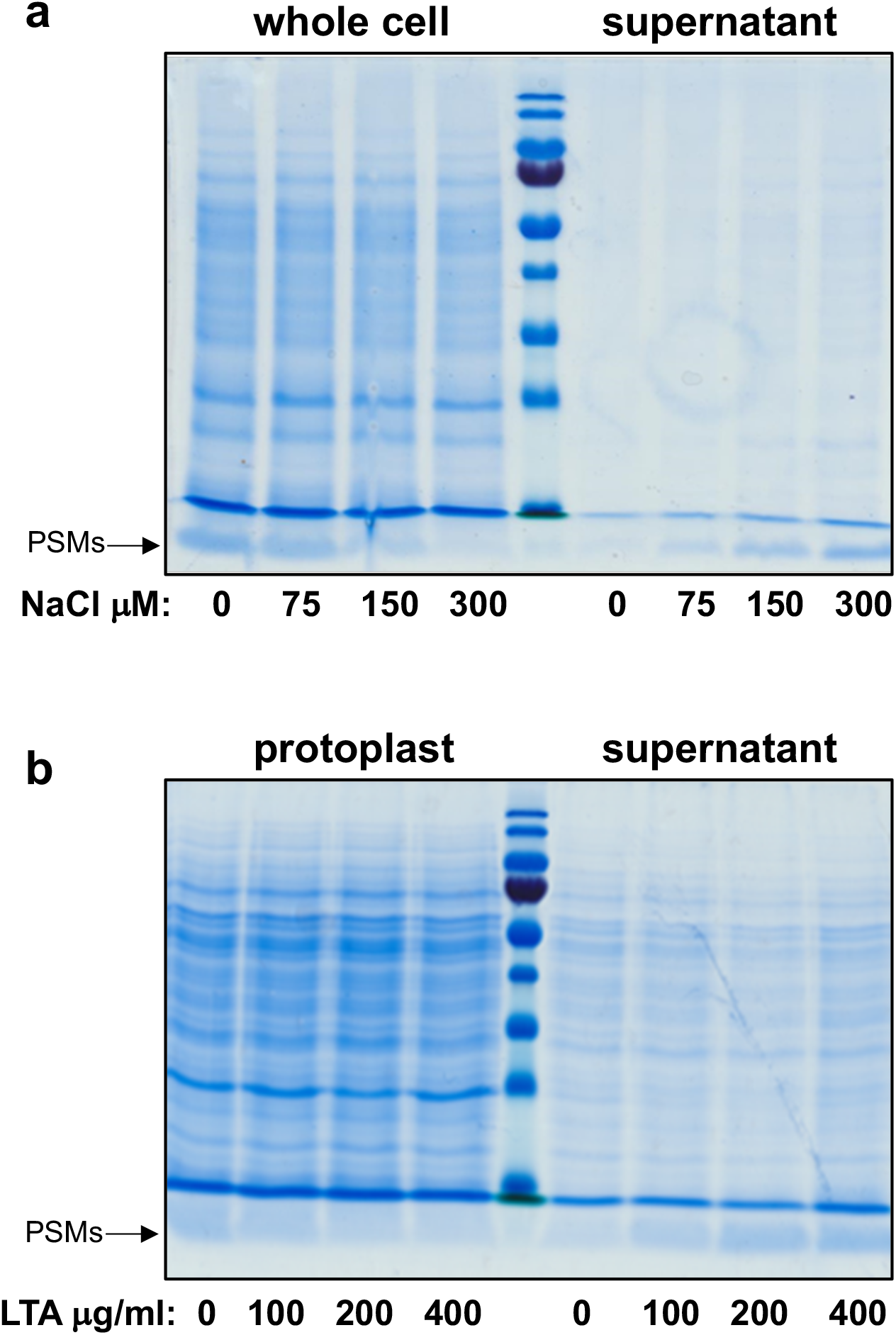
In the absence of WTA the PSMs are retained within the cells due to electrostatic interactions. (**a**) The PSMs of the TarO mutant moved from the cells into the supernatent when incubated in increasing concentrations of NaCl. (**b**) The PSMs of the TarO mutant moved from the protoplast into the supernatant when incubated with increased concentraions of LTA.

### WTA associated electrostatic charge across the cell wall affects the release of cytolytic toxins from the bacterial cell

The cell envelope of *S. aureus* is predominantly negatively charged, due to the presence of negatively charged molecules such as the phospholipids in the membrane, membrane-bound lipoteichoic acids (LTA) and cell wall-bound WTA. On the other hand, cytolytic toxins are predominantly positively charged. This led us to hypothesise that these toxins may not be released from WTA mutants’ cells due to the changes in electrostatic forces the lack of WTA causes, where the negatively charged WTA molecules may be involved in drawing the positively charged toxin away from the membrane and out towards the extracellular environment. To test this with a focus on the PSMs, we incubated TarO mutant cells in increasing concentrations of NaCl, where the increasing ionic strength should reduce any electrostatic interactions that may be occuring between the toxins and the negatively charged molecules within the membrane. This resulted in a decrease in the abundance of the PSM from the cells, and a corresponding increase in the levels of the PSMs in the supernatant, suggesting that in the absence of WTA, toxins are retained within the bacteria cells due to electrostatic interactions. We then speculated that toxins might be held within the cell envelope by interacting with LTA, so we tested whether purified soluble LTA could release PSMs from protoplasts. As with the increasing level of NaCl, with increasing concentrations of LTA we observed a reduction in the abundance of the PSMs associated with the protoplast, and an increase in their abundance released into the supernatant. This further strengthens our hypothesis that electrostatic interactions between the positively charged toxins and negatively charged molecules in the cell membrane are causing the retention of the toxins in the absence of WTA.

## Discussion

The production and release of cytolytic toxins is a polygentic trait that begins with the sensing by the bacteria of the external environment for conditions in which the production toxins will be most benefical, through a complex regulatory cascade that leads to the transcription of the toxin genes (37–40). Once translated the toxins are secreted through the cytoplasmic membrane, and up until now, their movement from the outer leaflet of the membrane was considered a passive diffusion process through the mesh like structure of peptidoglycan of the cell wall. However, in recent findings a role for specific macromolecules in the cell envelope in the release of cytolytic toxins have been described. Leukocidin AB (LukAB), has been found to be retained as discrete foci in two distinct compartments of *S. aureus* cells: membrane-proximal and surface-exposed, where lipoteichoic acid (LTA) was found contribute to this LukAB deposition and release (29). Further to this, here we demonstrate here that features such as electrostatic charge associated with key cell wall molecules such as WTA also play a critical role in the movement of toxin through the cell wall and their release into the extracellular environment. It is interesting to note that not all toxins were affected by the loss of WTA (Fig. 2a & b), why this is and how these other toxins traverse the cell wall is not as yet understood, demonstrating how much we have yet to learn about this critical pathogenic trait.

Several studies have demonstrated how WTA abundance and WTA modifications mediate staphylococcal virulence (41, 42). In particular, increased WTA production has been shown to promote abscesses formation, through an increased host immune response (41), while WTA glycosylation has been shown to have an impact on the adherence with host epithelium and to govern nasal colonisation (42). Despite this importance for survuval *in vivo*, the bacteria can survive *in vitro* without WTA so long as a gene upstream of the WTA biosynthetic pathway, such as TarO is inactivated (32, 33). Given its importance it is therefore interesting that we were unable to detect the transcription of the *tarF* gene using either a GFP fusion or qRT-PCR. It is possible that this is an *in vitro* effect, and that had we access to *in vivo* sampels we may have been able to detect transcription, and confirmed the effect the SNP identified amongst our clinical isoaltes (Fig. 1c) has on the transcription of *tarF*. An alternative explanation relates to the negative effect the inactivation other *tar* genes involved in the later stages of WTA biosynthesis have on the bacteria. Given the toxic effect to the bacteria either these gene products and/or partially formed WTA subunits have when imbalanced, it is likely that their expression would be tighly controlled. It is therefore possible what the window of time in which the transcription of these gene is detectable is limited, and missed by us under these experiment conditions. It could also be the case that these genes transcribed at a level too low to detect by the techniques used here, but understanding whether any of these explain why we don’t detect *tarF* transcript would require an indepth molecular investigation.

WTA is critical to the ability of *S. aureus* both colonise asymptomatically and to cause disease, and here, through the use of a functional genomic approach, we describe a further virulence mechanism regulated by WTA production and uncovered a hitherto unknown aspect to the cytotoxicity of a major human pathogen. Considered alongside other recent findings we propose that the movement of cytolytic toxins from the outer leaftet of the cytoplasmic membrane to the extracellular environment is a complex and largely opaque process involving the activity and physical presence of a number of cell wall associated macromolecules. It is a reseach area that warrants further detailed investigation, given the potential that the blocking of such processes could bring to the development of novel therapeutic strategies.

## Materials and methods

### Bacterial strains and growth conditions

The list of S. *aureus* strains is listed in Table 1. Tryptic soy broth (TSB) and Tryptic soy agar (TSA) were used for all *S. aureus* strains cultures. Strains containing transposon insertions were cultured on TSA plates with the addition of erythromycin (5μg/ml). Strains containing pRMC2 plasmid were selected on chloramphenicol (10 μg/ml), while anhydrous tetracycline (100 ng/μl or 200 ng/μl) was added to the media for the activation of the inducible promoter.

**Table 1.** List of strains used in this study.

| Strain name | Description | Reference |
| --- | --- | --- |
| LAC | USA300; CA -MRSA, type IV SCCmec | (46) |
| LAC $\Delta tarO$ | *tarO* KO mutant in LAC | (47) |
| 95E07 | Je2 strain with a transposon insertion in the *tarK tarF* intergenic region, Tn::298351 | (30) |
| LAC *tarKF::Tn* | LAC strain with transposon insertion in the *tarK tarF* intergenic region, Tn::298351 | This study |

**Table 2.**
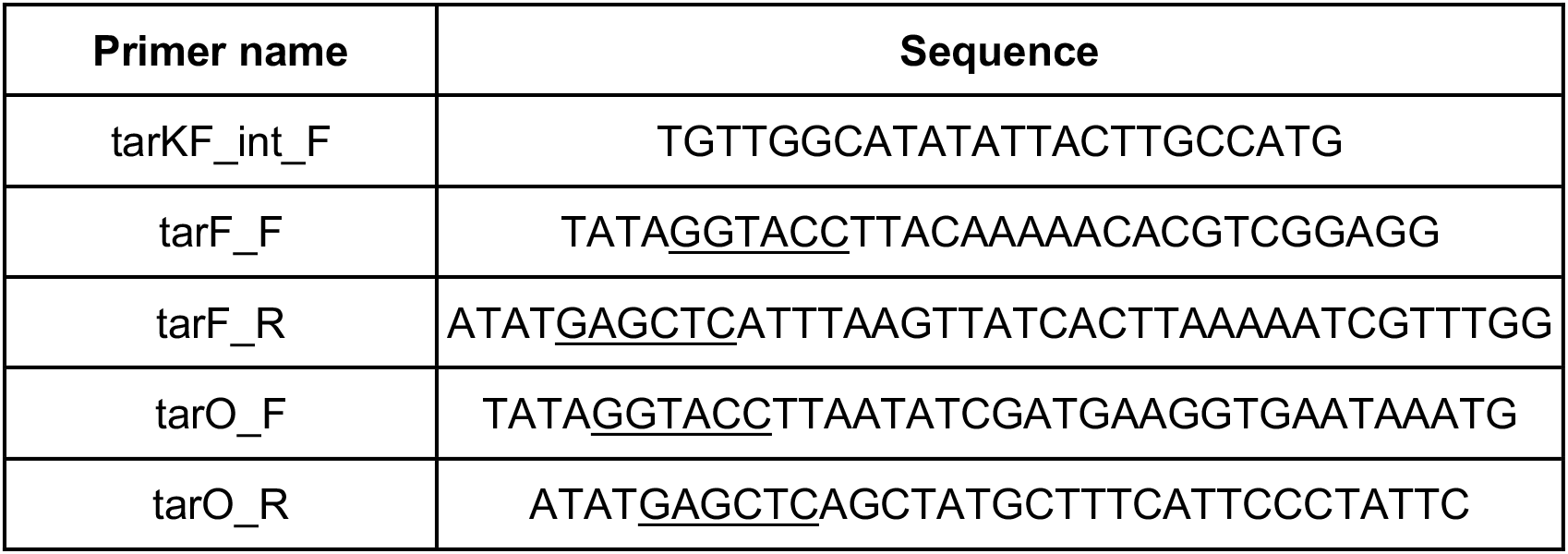
Primers used in this study.

### Bacterial genetic manipulation

The transposon insertion from the 95E07 strain (30) was transferred to LAC through phage transduction using phage Φ11, generating the strain LAC *tarKF::tn*. Correct transposon insertion was verified through colony PCR using the tarKF_int_F and tarF_R primers. For complementation experiments, the *tarF* gene was amplified using JE2 genomic DNA and KAPA HiFi polymerase (Roche) with the primers tarF_F and tarF_R, while the *tarO* gene was amplified using the primers tarO_F and tarO_R. tarF_F and tarO_F primers contained the KpnI restriction site at the 5’ end, while tarF_R and tarO_R included a SacI restriction site. The PCR products were cloned into the pRMC2 plasmid, using KpnI and SacI restriction enzymes and T4 ligase (NEB) to make the *ptarO* and *ptarF* plasmids. These were firstly transformed via electroporation to the RN4220 strain and eventually transformed into the LAC mutants to complement LAC Δ*tarO* and LAC *tarKF::tn* mutations.

### WTA extraction and analysis

WTA extraction was performed as described previously (43). Briefly, overnight cultures were centrifuged and washed with Buffer1 (50 mM MES, pH 6.5). The samples were boiled 1h in SDS containing Buffer2 (4% [wt/vol] SDS, 50 mM MES, pH 6.5) to remove all LTA and contaminating lipids. Samples were washed sequentially in Buffer1, Buffer2, Buffer3 (2% NaCl, 50 mM MES, pH 6.5) and Buffer1 to remove all lipids and residual SDS. Proteins were removed by digestion with proteinase K for 4h at 50°C, in digestion buffer (20 mM Tris-HCl pH 8.0, 0.5% [w/v] SDS). Samples were washed in Buffer3 and then three times in MilliQ H2O. Samples were then incubated 16h in 0.1 M NaOH, to release the WTA from the peptidoglycan. The samples were centrifuged, and the supernatants were neutralized with 1 M Tris-HCl, pH 7.8. The WTA extractions were stored at −20°C before proceeding with following analyses. Phosphorous amounts were measured from WTA extraction samples as previously described (44). WTA acrylamide gels were run as previously described (43) in Tris-Tricine buffer (1 M Tris, 1 M Tricine, pH 8.2). The gels were separated at 4°C and 40mA with constant stirring until the dye front reached the bottom. The gels were washed three times in MilliQ H2O, then stained overnight with Alcian blue solution (1 mg/ml Alcian blue, 3% acetic acid). Gels were then destained in MilliQ H2O until WTA became visible.

### THP-1 toxicity assay

The monocytic THP-1 cell line (ATCC TIB-202) was used as previously described (34, 45). Cells were cultivated in RPMI-1640 supplemented with heat-inactivated fetal bovine serum (10%), L-glutamine (1 M), penicillin (200 U/ml), and streptomycin (0.1 mg/ml) in a humidified incubator at 37°C with 5% CO2. For toxicity assays, cells were harvested by centrifugation and resuspended to a final density of 1 x 10^6^ to 1.5 10^6^ cells/ml in tissue-grade Hank’s balanced salt solution (HBSS), typically yielding 95% viability. Bacterial supernatants harvested from S. aureus overnight cultures (18h), were diluted to 10-30% in TSB and mixed 1:1 with THP-1s incubated for 12’ at 37°C. THP-1 cell death was measured by Trypan blue exclusion assay. Three technical replicates were performed for each biological replicate, independent biological replicates were performed in different days.

### Cell fractionation and protein analysis

Whole cell lysates were prepared from overnight bacterial cultures, normalized to same OD. Samples were centrifuged and the supernatant stored at −20°C. Cell pellets were washed once in PBS, then treated with lysostaphin to digest the cell wall. SDS was added to the suspensions at a final concentration of 2%, and the samples boiled for 20’. The samples were stored at - 20°C. Cell wall and protoplasts fractions were obtained from overnight cultures. Cell pellets were washed in TSM buffer (50 mM Tris-HCl pH 7.5, 0.5 M sucrose, 10 mM MgCl_2_), and treated with lysostaphin in TSM buffer for 15’ at 37°C. Samples were centrifuged at 8000rpm, and supernatants containing the cell wall fraction were stored at −20°C. The pellets, containing the protoplasts, were resuspended in PSB 2% SDS and boiled for 20’, then samples were stored at −20°C. For protein gels, samples were mixed with 2X SDS loading dye, boiled for 10’, and loaded onto 12% acrylamide gels. The gels were either Coomassie stained for PSMs and total protein visualised, or transferred to either a nitrocellulose or PVDF membrane for western blotting. Transfer was performed using either a Trans-Blot Turbo Transfer System (Bio-Rad), or an iBlot2 transfer device (Life Technologies). The membranes were blocked either overnight in PBS tween milk (10%) or for 10’ in StartingBlock T20 (TBS). Membranes were incubated with toxin specific antibodies (1:2000, 1μg/ml) in either PBS tween milk (3%) for 1h, or in StartingBlock T20 (TBS) overnight. Membranes were washed and incubated with horseradish peroxidase conjugated secondary antibodies for 1h. Proteins were detected by using either Opti-4CN detection kit (Bio-Rad) or ECL detection kit (Cytiva).

### PSMs band identification

Bacteria (JE2) were cultured for 18 h in 5 ml TSB and centrifuged to separate cell pellet from supernatant. Proteins in the supernatant fraction were concentrated via Trichloroacetic acid (TCA) precipitation. Briefly, 1 volume of cold (4°C) TCA (VWR) was mixed with 4 volumes of supernatant and incubated on ice for 60 minutes. Precipitated proteins were pelleted by centrifugation and washed in ice cold acetone three times. Protein pellets were resuspended in 50 uL 8 M Urea. Protein samples were then mixed with Blue Protein Loading Dye (NEB) boiled for 10 minutes and separated on 10% SDS-PAGE gel for 50 minutes at 140 V. The gel was stained with Quick Coomassie (Generon) for 2 hours and de-stained overnight in deionised water. The band below the dye-front, corresponding to less than 5 kDa was excised, stored in deionised water and delivered to the Bristol Proteomics Facility where the band was subject to in-gel proteolytic digestion using an automated DigestPro. Resulting peptides were analysed by MALDI-ToF MS/MS using a Bruker Daltonics UltrafleXtreme 2 Mass Spectrometer.

### Dextran diffusion assay

Murein sacculi were extracted following the same protocol of WTA extraction, with the exclusion of the final NaOH treatment. Sacculi were incubated overnight in a solution of 5μM 40 kDa dextran labelled with Texas red. Samples were diluted 1:50 in water, and the fluorescence of the sacculi was followed by FACS for 30’. Mean fluorescence of 1 minute intervals were used to compare different samples.

### Release of toxins by NaCl and LTA

Cells from overnight cultures were pelleted by centrifugation, and subsequently washed twice in PBS. The samples were resuspended in PBS with increasing concentrations of NaCl, and incubated at 37°C, 180 rpm for 1h. The cells suspensions were then centrifuged, the supernatants stored at −20°C. The pellets, containing the cells, were treated as described previously for whole cell lysates preparation. Protoplasts were isolated as described previously for cell fractionation. The protoplasts were resuspended in TSM with increasing concentration of LTA and incubated at 37°C 180 rpm shaking for 10 minutes. The samples were then centrifuged for 5’ at 5000 rpm, supernatants were stored at −20°C, while pellets were resuspended in TSM buffer 2% SDS.

### Electron Microscopy

Cells from overnight cultures were pelleted by centrifugation, washed once in PBS and resuspended in 2.5% glutaraldehyde in 0.1 M sodium cacodylate buffer, pH 7.2. Fixed bacteria were delivered to the Wolfson Bioimaging Facility, University of Bristol where they were dehydrated, embedded in resin, stained, and sectioned. Images were acquired using a FEI Tecnai 12 electron microscope.

### Statistical analysis

Statististical analyses were performed using GraphPad Prism 8.4.

## Acknowledgements

We would like to thank Simon Foster, Laia Pasquina Lemonche and Richard Daniel for helpful discussion of the data. We would also like to acknowledge the assistance of Dr. Andrew Herman and Helen Rice for cell sorting and the University of Bristol Faculty of Life Sciences Flow Cytometry Facility, and Chris Neal, Mark Jepson and Lorna Hodgson from the Wolfson Bioimaging Facility for their electron microscopy services.

